# Scalp EEG interictal high frequency oscillations as an objective EEG biomarker of infantile spasms

**DOI:** 10.1101/2020.05.31.126573

**Authors:** Hiroki Nariai, Shaun A. Hussain, Danilo Bernardo, Hirotaka Motoi, Masaki Sonoda, Naoto Kuroda, Eishi Asano, Jimmy C. Nguyen, David Elashoff, Raman Sankar, Anatol Bragin, Richard J. Staba, Joyce Y. Wu

**Author notes:** Corresponding Author: Hiroki Nariai, MD, PhD, Division of Pediatric Neurology, Department of Pediatrics, UCLA Mattel Children’s Hospital Address: 10833 Le Conte Ave, Room 22-474, Los Angeles, CA 90095-1752, U.S.A.

## Abstract

**Objective:** To investigate the diagnostic utility of high frequency oscillations (HFOs) via scalp electroencephalogram (EEG) in infantile spasms.

**Methods:** We retrospectively analyzed interictal slow-wave sleep EEGs sampled at 2,000 Hz recorded from 30 consecutive patients who were suspected of having infantile spasms. We measured the rate of HFOs (80-500 Hz) and the strength of the cross-frequency coupling between HFOs and slow-wave activity (SWA) at 3-4 Hz and 0.5-1 Hz as quantified with modulation indices (MIs).

**Results:** Twenty-three patients (77%) exhibited active spasms during the overnight EEG recording. Although the HFOs were detected in all children, increased HFO rate and MIs correlated with the presence of active spasms (p < 0.001 by HFO rate; p < 0.01 by MIs at 3-4 Hz; p = 0.02 by MIs at 0.5-1 Hz). The presence of active spasms was predicted by the logistic regression models incorporating HFO-related metrics (AUC: 0.80-0.98) better than that incorporating hypsarrhythmia (AUC: 0.61). The predictive performance of the best model remained favorable (87.5% accuracy) after a cross-validation procedure.

**Conclusions:** Increased rate of HFOs and coupling between HFOs and SWA are associated with active epileptic spasms.

**Significance:** Scalp-recorded HFOs may serve as an objective EEG biomarker for active epileptic spasms.

**Highlights:**

- Objective analyses of scalp high frequency oscillations and its coupling with slow-wave activity in infantile spasms were feasible.
- Increased rate of high frequency oscillations and its coupling with slow-wave activity correlated with active epileptic spasms.
- The scalp high frequency oscillations were also detected in neurologically normal children (although at the low rate).

## 1. INTRODUCTION

Infantile spasms is an often devastating epileptic syndrome in infancy, with an incidence ranging from 2 to 3.5 per 10,000 live births (Pellock et al., 2010). It is characterized by epileptic spasms (frequently in clusters), distinctive interictal electroencephalography (EEG) patterns (hypsarrhythmia), and arrest or regression in psychomotor development regardless of the etiology (Lux and Osborne, 2004; Osborne et al., 2019; Pellock et al., 2010; Wirrell et al., 2015). Lead time to treatment is a major factor affecting future neurodevelopmental outcomes, and the earlier treatment is believed to result in better outcomes (Koo et al., 1993; O’Callaghan et al., 2011). Thus, prompt diagnosis and treatment (with adrenocorticotropic hormone (ACTH), high-dose oral prednisolone, and/or vigabatrin) are crucial in managing children with infantile spasms. Scalp EEG is an essential clinical test to aid the early diagnosis. The presence of hypsarrhythmia during interictal background activity is a specific finding of infantile spasms. In spite of hypsarrhythmia’s seemingly simple definition (interictal EEG patterns with high voltage, disorganization, and multifocal independent epileptiform discharges) (Lux and Osborne, 2004; Noachtar et al., 1999), the definition is imprecise and is open to subjective interpretation that contributes to low inter-rater reliability even among experienced pediatric electroencephalographers (Hussain et al., 2015). Having more objective, accurate interictal EEG features for the detection of active infantile spasms is desperately needed.

High frequency oscillations (HFOs) (80-500 Hz) recorded with invasive intracranial EEG have been reported as a promising biomarker of epileptogenic brain regions in both adults and children (Akiyama et al., 2011; Bragin et al., 1999a; Bragin et al., 1999b; Jacobs et al., 2010; Jirsch et al., 2006; Nariai et al., 2011b; Staba et al., 2002; Urrestarazu et al., 2007; Worrell et al., 2004; Wu et al., 2010). Recent studies have shown EEG fast activities can be recorded non-invasively from the scalp recordings (Andrade-Valenca et al., 2011; Bernardo et al., 2018; Kobayashi et al., 2004; Kobayashi et al., 2015; Kramer et al., 2019; Nariai et al., 2017; Nariai et al., 2018; Pizzo et al., 2016; Wu et al., 2008). Scalp EEG HFOs have the potential to localize seizure onset zones (Andrade-Valenca et al., 2011; Bernardo et al., 2018; Nariai et al., 2017; Tamilia et al., 2020), and to monitor disease activity in epileptic spasms (Kobayashi et al., 2015), Rolandic epilepsy (Kobayashi et al., 2011; van Klink et al., 2016), and epileptic encephalopathy with continuous spike-and-wave during sleep (CSWS) (Cao et al., 2019; Kobayashi et al., 2010). Additionally, HFOs coupled with slow-wave activity strongly localize to epileptic brain regions in children with infantile spasms and other types of focal epilepsy in invasive monitoring (Iimura et al., 2018; Motoi et al., 2018; Nonoda et al., 2016; Song et al., 2017; Weiss et al., 2016) and may reflect disease activity in scalp EEG in epileptic spasms (Bernardo et al., 2020. In press; Iimura et al., 2019). Specifically, HFOs and SWA bands at 3-4 Hz and 0.5-1 Hz are reportedly coupled with pathological HFOs in children with infantile spasms (Iimura et al., 2018; Nariai et al., 2011a).

The objective of this study was to quantify the rate of HFOs and the degree of cross-frequency coupling of HFOs and slow-wave activity on scalp EEG in children who presented with a clinical suspicion of infantile spasms. We then investigated whether such objective HFO-related metrics would accurately predict the presence or absence of epileptic spasms.

## 2. METHODS

### 2.1. Cohort

This is a retrospective cohort study of infants who were suspected of having infantile spasms based on the clinical history and underwent overnight video-EEG evaluation at UCLA between October 2016 and May 2019. Inclusion criteria consisted of (1) age ≤ 12 months, (2) availability of at least one 10-minute epoch of slow-wave sleep EEG sampled at 2,000 Hz, and (3) clinical follow-up for at least one month. EEG recordings with more than two channels with significant artifacts per recording were excluded from the study (the determination was made before the quantitative EEG analysis). There were no other exclusion criteria.

### 2.2. Standard protocol approval

The institutional review board at UCLA approved the use of human subjects and waived the requirement for informed consent.

### 2.3. Electroencephalography (EEG) recording

Scalp EEG recordings were obtained using Nihon Kohden (Irvine, California) EEG acquisition systems, utilizing 21 gold-plated electrodes placed according to the international 10–20 system, and a digital sampling frequency of 2000 Hz. The EEG recording defaults to a proprietary setting of a low frequency filter of 0.016 Hz and a high frequency filter of 600 Hz at the time of acquisition. The study subjects were identified by DB and HN, and an artifact-free 10-minute epoch of non-REM sleep EEG devoid of the obvious artifact was selected by a board-certified pediatric electroencephalographer (HN) while blinded to the clinical information. A longitudinal bipolar montage was used for the quantitative EEG analysis.

### 2.4. Characterization of epileptic spasms and hypsarrhythmia

As a part of routine clinical evaluation, a board-certified pediatric electroencephalographer determined (1) the presence or absence of epileptic spasms, and (2) the presence or absence of hypsarrhythmia, defined as EEG patterns with high voltage (non-epileptiform slow-waves with amplitude consistently greater than 200 μV), disorganization (loss of the anterior-posterior amplitude-frequency gradient), and abundant multifocal independent epileptiform discharges (Hussain et al., 2015). Patients with epileptic spasms were treated, and on the basis of routine follow-up video-EEG, they were either classified as “responders” (i.e. freedom from epileptic spasms and hypsarrhythmia, without relapse for at least one month) or “non-responders” (persistent epileptic spasms and/or hypsarrhythmia).

### 2.5. Computational EEG analyses

Computational analyses were accomplished using a laptop PC equipped with an Intel Core processor (i5-7200U, 2.50 GHz), 8 GB random access memory (RAM), Microsoft Windows (version 10, 64 bit), and MATLAB software (version 2018b, MathWorks, MA). We specifically calculated the rate of HFOs and cross-frequency coupling of HFOs and slow-wave activity (SWA).

#### 2.5.1 Semi-automated analysis of interictal scalp HFOs

We defined interictal high frequency oscillations (HFOs) as oscillatory events with at least six cycles, a center frequency occurring between 80-500 Hz, which stands out from the background signal. Using a 10-minute epoch of EEG during slow-wave sleep, without epileptic spasms during the period, HFOs were preliminarily identified by an automated short-term energy detector (RippleLab; MATLAB-based open-source program) (Navarrete et al., 2016; Staba et al., 2002). Both the RMS standard deviation (SD) threshold and the peak SD threshold of putative HFO events were set at 3 SD. We then visually confirmed HFOs by excluding false-positive events, namely ringing (filtering of sharp transients)(Benar et al., 2010), as well as muscle and electrode artifacts (see examples in **Figure 1**). The detected events before the visual verification (automated analysis only), and the events after the visual verification (semi-automated analysis with visual verification) were used for further correlative analysis. Also, we separated HFOs as ripples (80-250 Hz) and fast ripples (FR: 250-500 Hz) to assess whether all HFOs or a subset of HFOs could predict epileptic spasms. The number of verified events were recorded in each of 20 channels (Fp2-F8, F8-T4, T4-T6, T6-O2, Fp1-F7, F7-T3, T3-T5, T5-O1, T4-C4, C4-Cz, Cz-C3, C3-T3, Fp2-F4, F4-C4, C4-P4, P4-O2, Fp1-F3, F3-C3, C3-P3, P3-O1). For each subject, the HFOs rate (events per minute) was defined as the average rate across all 20 channels (See example in **Figure 2**).

**Figure 1:**
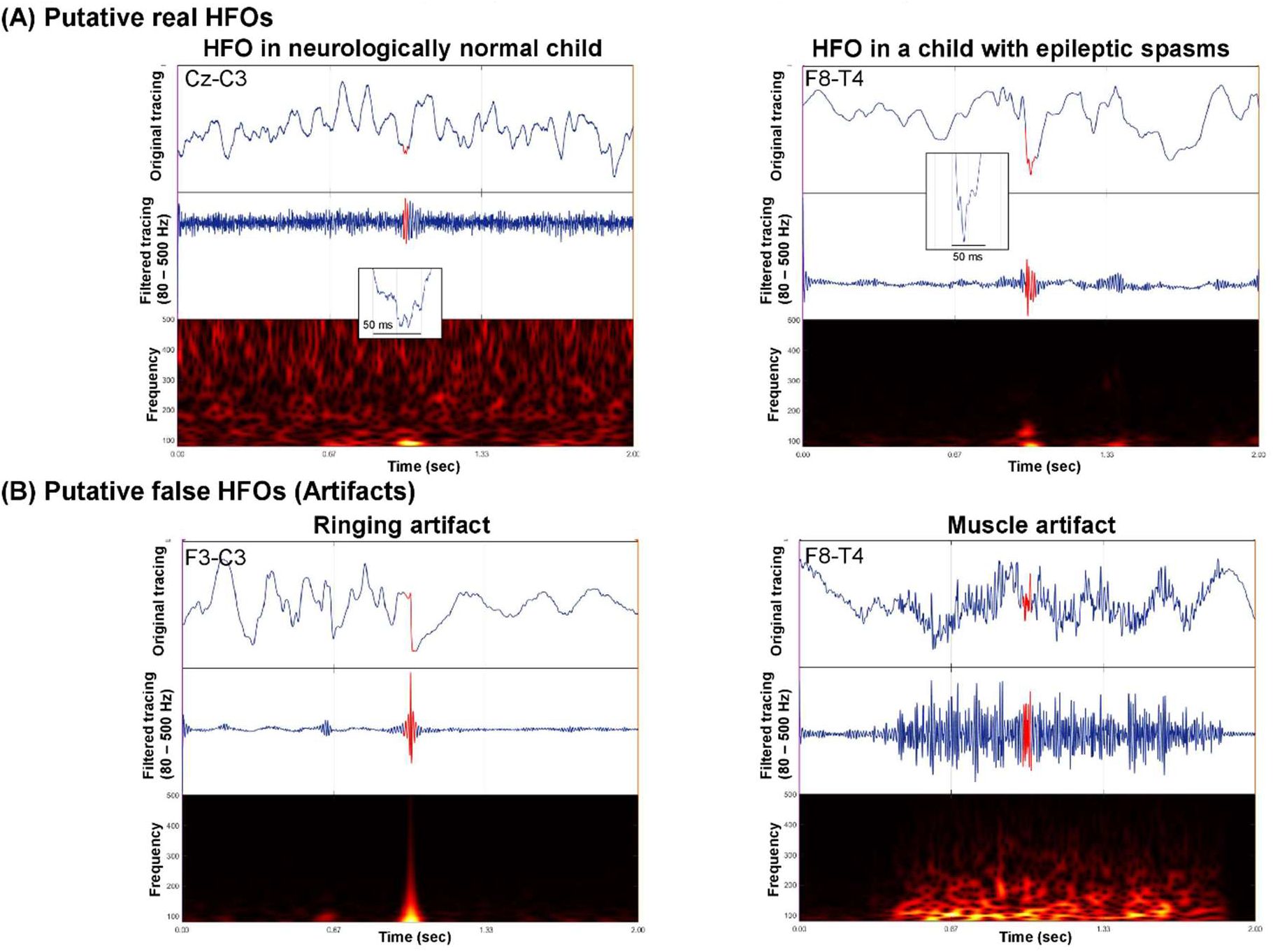
HFO detection (true positives and false positives) Examples of putative real HFOs (A) and false HFOs (B) in automated HFOs detection using RippleLab are shown. (A) Time-frequency plots of HFO in a neurologically normal child (left, patient #4) and HFO in a child with epileptic spasms (right, patient#23) are shown. (B) Time-frequency plots of ringing artifact (left) and muscle artifact (right) are shown.

**Figure 2:**
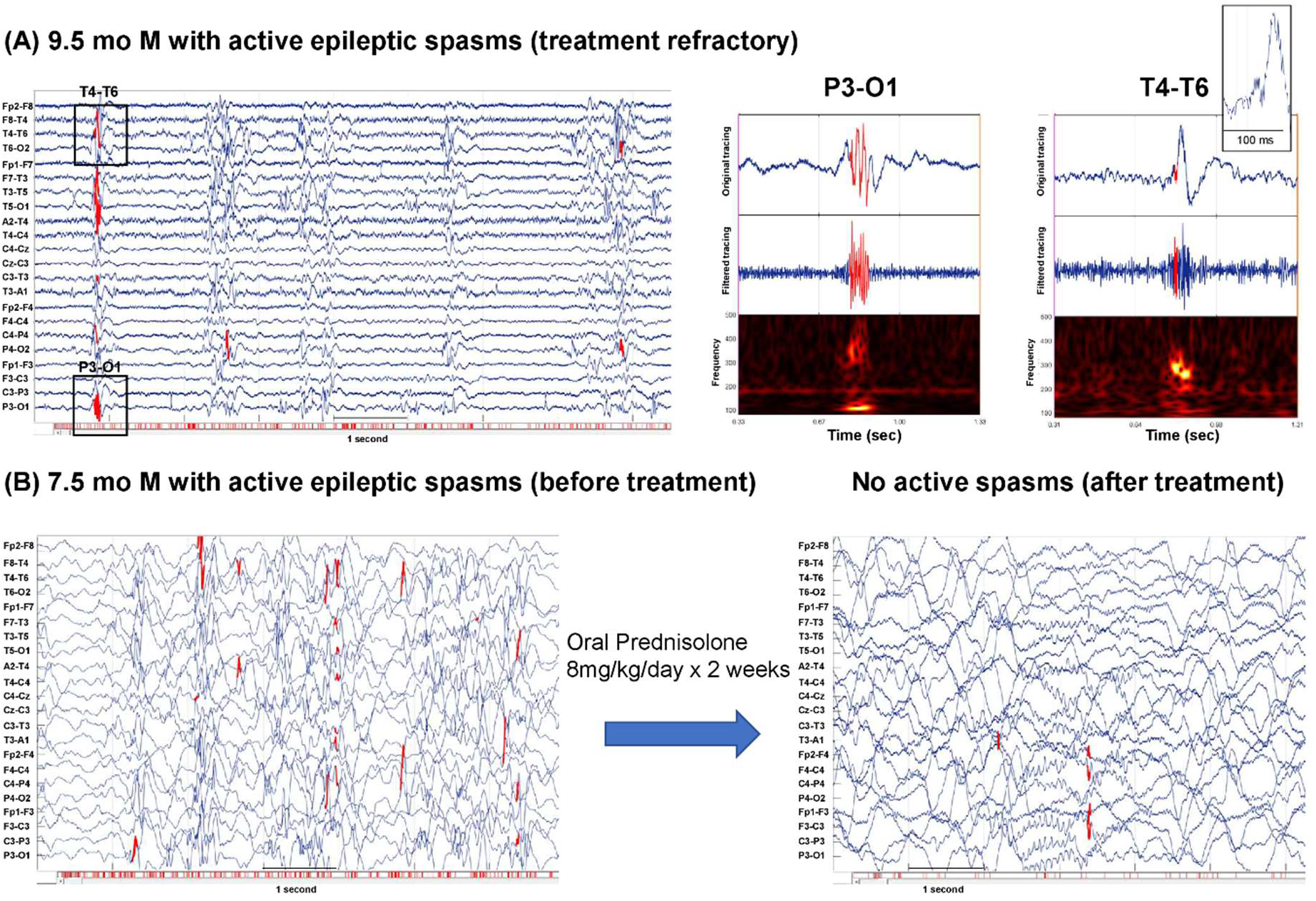
HFO detection and clinical outcome. (A) Scalp EEG tracing in the bipolar montage is shown (left). Red markings within the EEG tracings show detected HFO events. The bottom horizontal bar shows detected HFO events during the entire time period (10 minutes). Examples of time-frequency plots are shown (right). HFOs in the ripple band (80-250 Hz) at P3-O1 and the fast ripple band (250-500 Hz) at T4-T6 are shown. (B) Scalp EEG tracings in the bipolar montage before (left) and after (right) high-dose prednisolone treatment are shown. Red markings show detected HFO events. The bottom horizontal bar shows detected HFO events during the entire time period (10 minutes each). A significant decrease in HFO rate after the treatment is noted.

#### 2.5.2. Fully automated analysis of cross-frequency coupling of HFOs and slow-wave activity (SWA)

The modulation index (MI) represents the strength of the coupling between fast oscillations amplitude and slow-wave activity (SWA) phase (Canolty et al., 2006; Tort et al., 2010). The EEGLab, PACT plugin (MATLAB-based open-source program) was used to compute the MIs in each channel similar to previous studies (Iimura et al., 2018; Miyakoshi et al., 2013; Nonoda et al., 2016). With the assumption that pathological HFOs are often high-amplitude and coupled with SWA activity (Nonoda et al., 2016; Song et al., 2017), the MI was calculated using the subset of automatically-detected HFOs, the amplitude of which was greater than the 98^th^ percentile (within-subject). Thus, no visual verification for HFO detection was performed in the calculation. We specifically calculated two MIs (HFOs & 3-4 Hz and HFOs & 0.5-1 Hz) in each channel, as both SWA bands at 3-4 Hz and 0.5-1 Hz are reportedly coupled with pathological HFOs in children with infantile spasms (Iimura et al., 2018; Nariai et al., 2011a). The MIs for each subject were defined as the mean MI across of all channels, similar to our approach for HFO rate, above. (See analysis example in **Figure 3**.)

**Figure 3:**
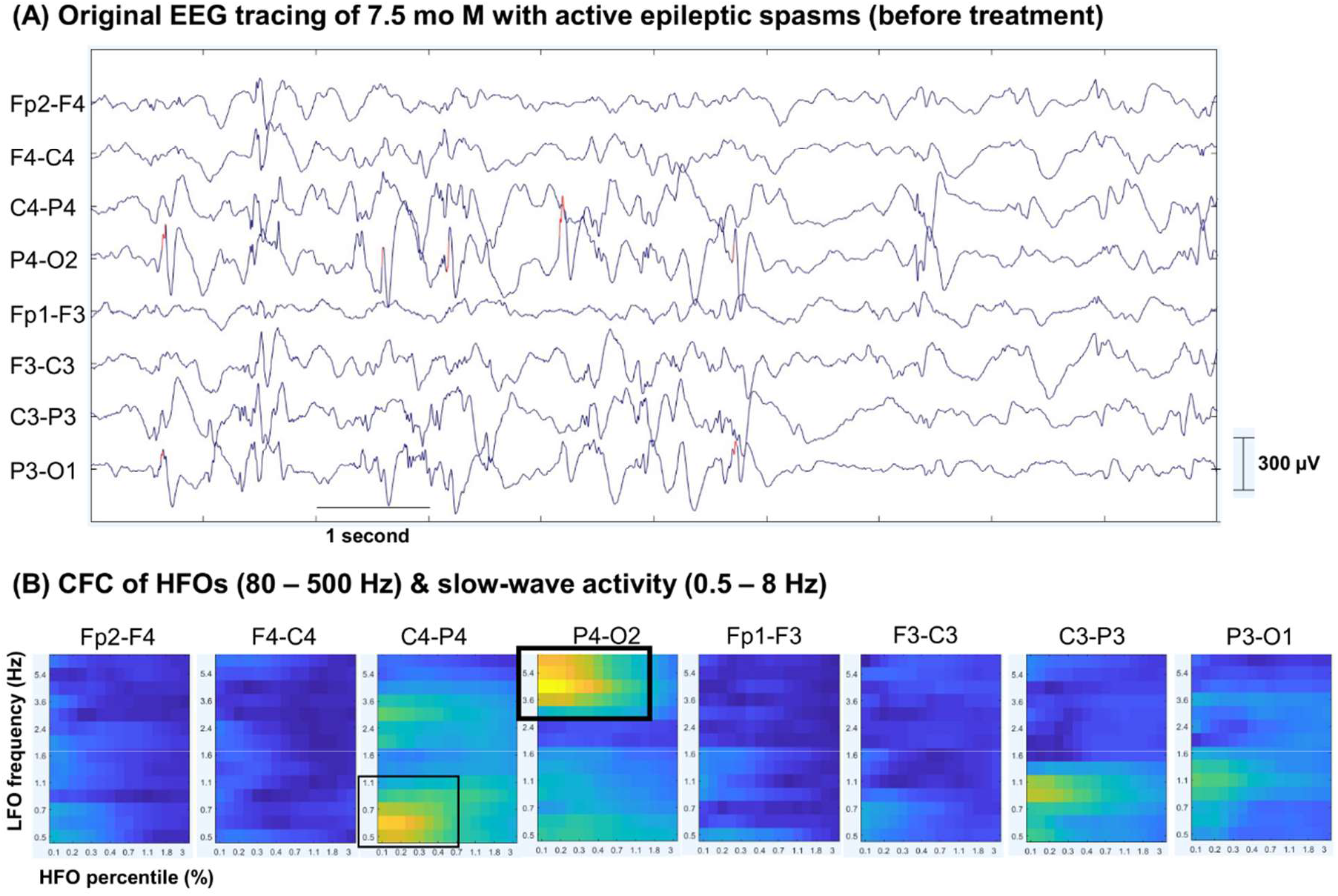
Cross-frequency coupling of HFOs & slow-wave activity. (A) Scalp EEG tracing using the longitudinal bipolar montage is shown. Red markings show detected HFOs (80-500 Hz) events for cross-frequency coupling (CFC) analysis. (B) Heat-maps of the CFC analysis in each channel are shown. The x-axis shows the HFOs (80-500 Hz) amplitude percentile (%), and Y-axis shows the low-frequency oscillation (LFO: 0.5-8 Hz) frequency. The yellow color indicates high modulation index (MI), while the dark blue color indicates low MI. High MI values of HFOs (80-500 Hz) & 3-4 Hz at channel P4-O2 is noted (thick black square). Mild to medial range MI values at 0.5-1 Hz at channel C4-P4 was also noted (thin black square).

### 2.6. Asymmetry in scalp EEG HFOs in relation to neuroimaging findings

Based on the HFO-related values, we calculated the asymmetry index (AI: [R - L] / [R + L]). The values range from −1 (values on the left hemisphere being disproportionally large) to +1 (values on the right hemisphere being disproportionally large) (Nariai et al., 2017). We compared the absolute asymmetric indices (abs AI) of HFOs (80-500 Hz) rate, MI of HFOs (80-500 Hz) & SWA (3-4 Hz and 0.5-1 Hz) between subjects with and without focal lesion on MRI.

### 2.7. Statistical analysis

Statistical results were obtained with JMP Pro (version 14; SAS Institute, U.S.A). The Wilcoxon rank-sum and Fisher exact tests were used to compare continuous and categorical variables, respectively. The Wilcoxon signed rank-sum test was used to compare matched pair values. For each EEG-related metric (HFO rate, MI of HFOs & 3-4 Hz, MI of HFOs & 0.5-1 Hz, and the presence of hypsarrhythmia), logistic regression models were created, and we compared the predictive model performances (the presence of epileptic spasms as the outcome) using area under the receiver-operator characteristic (ROC) curves. Based on the small number of subjects, we did not create multiple logistic regression models.

## 3. RESULTS

### 3.1. Patient characteristics

We identified 34 children, excluded 4 subjects due to significant artifacts in EEG recording, and included 30 children (15 females, 15 males) with a median age of 6.4 months (range: 1.7 – 11.0 months) for further quantitative analysis. Twenty-three (77%) exhibited epileptic spasms during the overnight recording. Of the 7 children without recorded epileptic spasms, one had a prior history of epileptic spasms (with abnormal EEG background). The remaining six children exhibited non-epileptic events only (i.e., normal infant behaviors and gastroesophageal reflux). They were neurologically normal and had no brain-based diagnosis such as epilepsy, autism, or developmental delay. Of the 23 children with epileptic spasms, 8 children had serial EEG recordings (two studies per subject) following treatment for epileptic spasms. Two out of 8 children were classified as responders and the rest were classified as non-responders. Details of patient characteristics were listed in **Table 1**.

**Table 1.** Patient characteristics and analysis results.

| Pt. No | Sex | Serial EEGs | Age at EEG (mo) | Diagnosis of IS | Active spasms on EEG | Timing of EEG | Response to Treatment | Treatment given | Etiology of IS | Neuroimaging | Hypser-rhythmia | Mean HFOs (80-500 Hz) rate (/min) | MI HFOs & 0.5-1 Hz | MI HFOs & 3-4 Hz |
| --- | --- | --- | --- | --- | --- | --- | --- | --- | --- | --- | --- | --- | --- | --- |
| 1 | M | N | 3.1 | N | N | Initial | NA | NA | NA | NA | N | 0.20 | 0.07 | 0.04 |
| 2 | F | N | 3.0 | N | N | Initial | NA | NA | NA | NA | N | 0.03 | 1.92 | 0.54 |
| 3 | F | N | 3.7 | N | N | Initial | NA | NA | NA | NA | N | 0.06 | 0.13 | 0.10 |
| 4 | F | N | 4.2 | N | N | Initial | NA | NA | NA | NA | N | 0.24 | 0.15 | 0.16 |
| 5 | F | N | 5.3 | N | N | Initial | NA | NA | NA | NA | N | 0.33 | 0.07 | 0.08 |
| 6 | F | N | 5.7 | N | N | Initial | NA | NA | NA | NA | N | 0.05 | 0.24 | 0.06 |
| 7 | F | N | 6.5 | Y | N | Repeat | Responder | TPM, VGB, ACTH, CBZ | Known (22q11.21 microduplication) | Periventricular leukomalacia with thinning of the corpus callosum | N | 0.13 | 0.07 | 0.07 |
| 8 | M | N | 6.1 | Y | Y | Initial | Responder | PRED, VGB | Known (severe HIE) | Diffuse cerebral injury consistent with HIE | N | 0.31 | 0.17 | 0.10 |
| 9 | M | N | 7.0 | Y | Y | Repeat | Responder | PRED, ACTH, VGB | Known (Lissencephaly) | Diffuse lissencephaly | N | 0.74 | 0.09 | 0.07 |
| 10 | M | N | 8.1 | Y | Y | Repeat | Non-responder | PRED, ACTH, VGB | Unknown | Diffuse cerebral volume loss | N | 1.25 | 0.23 | 0.08 |
| 11 | F | N | 9.5 | Y | Y | Repeat | Non-responder | PRED, ACTH, VGB | Unknown | Normal | N | 3.30 | 0.86 | 0.84 |
| 12 | F | N | 8.9 | Y | Y | Repeat | Non-responder | ACTH, VGB, CLB, ZNS | Known (CDKL5) | Left hippocampal malrotation | Y | 4.77 | 3.58 | 1.62 |
| 13 | M | N | 3.7 | Y | Y | Repeat | Non-responder | PB, LVT, PRED | Known (hemimegalencephaly) | Left hemimegalencephaly (s/p left hemispherectomy) | N | 4.72 | 1.44 | 1.51 |
| 14 | M | N | 8.5 | Y | Y | Repeat | Non-responder | PB, LVT, ACTH | Known (Lissencephaly) | Diffuse lissencephaly | N | 1.16 | 0.27 | 0.16 |
| 15 | M | N | 3.4 | Y | Y | Repeat | Non-responder | LVT, TPM, PB | Known (non-ketotic hyperglycinemia) | Normal | N | 1.66 | 0.70 | 0.20 |
| 16 | F | N | 6.4 | Y | Y | Repeat | Non-responder | B6, PRED, ACTH, VGB | Unknown | NA | N | 8.64 | 1.14 | 0.91 |
| 17 | F | N | 9.4 | Y | Y | Repeat | Responder | PRED, ACTH, VGB | Known (FCD) | Left posterior parietal FCD | N | 6.74 | 3.10 | 2.29 |
| 18 | M | N | 11.0 | Y | Y | Repeat | Non-responder | PRED, VGB, VPA | Known (Menkes disease with ATP7A mutation) | Bilateral subdural fluid collection | N | 1.04 | 0.11 | 0.11 |
| 19 | F | N | 4.5 | Y | Y | Initial | Responder | PRED, VGB | Unknown | Normal | N | 6.95 | 2.46 | 1.44 |
| 20 | F | N | 5.8 | Y | Y | Initial | Responder | PRED, VGB | Known (FCD) | Right occipital FCD | N | 4.84 | 0.73 | 0.56 |
| 21 | F | N | 1.7 | Y | Y | Initial | Non-responder | LVT, TPM, VGB | Known (Aicardi syndrome) | Agenesis of the corpus callosum, abnormal cortical gyration | N | 1.87 | 0.24 | 0.23 |
| 22 | M | N | 6.3 | Y | Y | Initial | Responder | PRED, VGB | Known (Trisomy 21) | Normal | Y | 5.55 | 2.32 | 0.98 |
| 23 | M | Y | 7.6 [8.1] | Y | Y [N] | Initial | Responder | PRED, VGB, ACTH | Unknown | Normal | N [N] | 6.37 [0.48] | 2.13 [0.12] | 1.23 [0.11] |
| 24 | F | Y | 5.6 [6.0] | Y | Y [N] | Initial | Responder | VGB | Known (TSC) | Numerous cortical/subcortical tubers (more prominent on the right hemisphere) and subependymal nodules | N [N] | 2.03 [0.47] | 0.80 [0.17] | 0.26 [0.10] |
| 25 | M | Y | 10.0 [13.7] | Y | Y [Y] | Repeat | Non-responder | VGB, ACTH, LVT, CLB | Known (FCD) | Right temporal parietal occipital FCD | N [N] | 2.41 [0.62] | 0.98 [0.44] | 0.79 [0.12] |
| 26 | M | Y | 6.1 [6.6] | Y | Y [Y] | Repeat | Non-responder | PRED, ACTH, VGB | Known (Ohtahara syndrome) | Moderate generalized volume loss | Y [Y] | 0.14 [2.21] | 0.12 [0.91] | 0.07 [0.43] |
| 27 | F | Y | 6.7 [9.1] | Y | Y [Y] | Repeat | Non-responder | PRED, ACTH, VGB | Known (ALG13 gene mutation) | Normal | N [Y] | 0.51 [3.43] | 0.12 [1.87] | 0.09 [1.75] |
| 28 | M | Y | 8.0 [8.5] | Y | Y [Y] | Repeat | Non-responder | ACTH, VGB | Known (BRAF gene mutation) | Generalized volume loss | Y [Y] | 6.23 [5.03] | 0.94 [1.60] | 0.65 [0.77] |
| 29 | M | Y | 7.2 [7.9] | Y | Y [Y] | Repeat | Non-responder | PB, LVT, CLB, VGB | Known (hemimegalencephaly) | Right hemimegalencephaly | Y [Y] | 6.14 [1.17] | 0.78 [0.09] | 0.75 [0.12] |
| 30 | M | Y | 7.9 [8.3] | Y | Y [Y] | Repeat | Non-responder | PRED, ACTH, VGB | Unknown | Normal | N [N] | 0.95 [0.19] | 0.49 [0.15] | 0.46 [0.10] |

IS: Infantile spasms; MI: Modulation index; HFO: High frequency oscillations; NA: Not applicable; FCD: Focal cortical dysplasia; TSC: Tuberous sclerosis complex; PRED: Prednisolone; TPM: Topiramate; VGB: Vigabatrin; ACTH: Adrenocorticotrophic hormone; CBZ: Clobazam; LTV: Levetiracetam; ZNS: Zonisamide; PB: Phenobarbital; VPA: Valproate; B6: Vitamine B6 [ ] indicates repeat values

### 3.2. Interictal scalp HFOs rate

HFO-related metrics in relation to the presence or absence of epileptic spasms are summarized in **Figure 4**. The HFO (80-500 Hz) rate after visual verification was higher among subjects with epileptic spasms (median: 2.41/min, IQR: 1.04, 6.14) than children without epileptic spasms (median: 0.13/min, IQR: 0.05, 0.24), with p < 0.001. Candidate HFOs (80-500 Hz) were detected in all subjects and the median HFO rate per channel before visual verification was 1.76/min (IQR: 0.51, 5.77). After the visual inspection, 13% of automatically-detected events were rejected, and authentic HFOs were recorded in all subjects, including in neurologically normal children. The events detected in the temporal channels (Fp2-F8, F8-T4, T4-T6, Fp1-F7, F7-T3, T3-T5) were rejected more often, compared to that in the parasagittal channels (T6-O2, T5-O1, C4-Cz, Cz-C3, F4-C4, C4-P4, P4-O2, F3-C3, C3-P3, P3-O1) (median removal rate/min: 0.27 [IQR: 0.09, 0.80] vs. 0.10 [IQR: 0.02, 0.25], p < 0.001). The median analysis time required for automated detection of HFOs (80-500 Hz) per study was 6.7 minutes (IQR: 5.9, 7.1), and the median time required to complete visual verification was 32.9 minutes (IQR: 27.5, 45.2) per subject.

**Figure 4:**
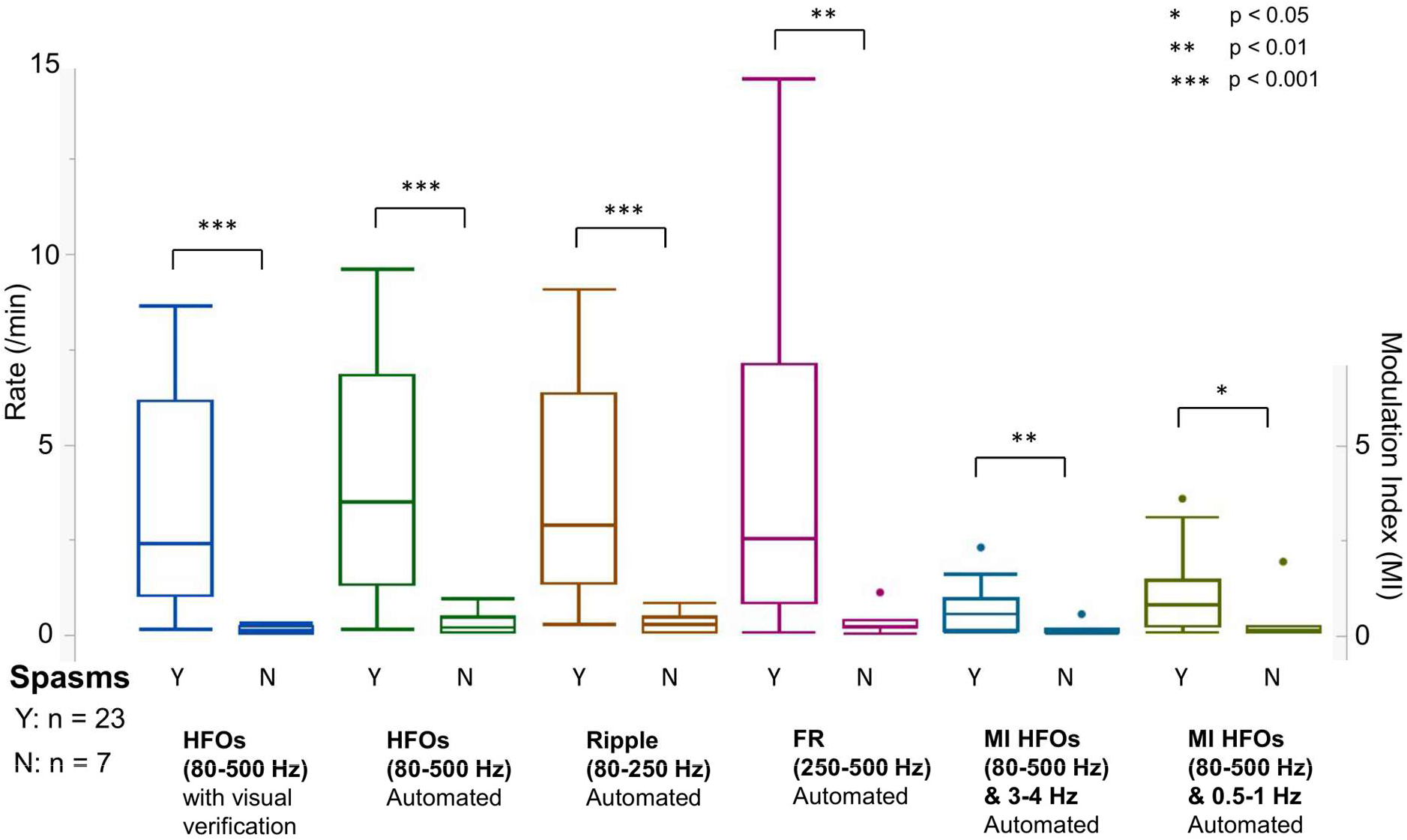
HFOs and epileptic spasms. HFO-related values (HFOs with or without visual verification, ripple, FR, MI of HFOs & 3-4 Hz and 0.5-1 Hz) are plotted as a function of the presence (Y) or absence (N) of the epileptic spasms. Note all the HFO-related values are higher in a group with active epileptic spasms.

### 3.3. Cross-frequency coupling of HFOs and slow-wave activity (SWA)

MIs of HFOs and 3-4 Hz SWA and HFOs and 0.5-1 Hz SWA were higher in children with than those without epileptic spasms (p < 0.01 and p = 0.02) (**Figure 4**). The median analysis time for modulation indices (MIs) of HFOs (80-500 Hz) & 3-4 Hz per case was 0.6 minutes (IQR: 0.55, 0.65) and that for HFOs (80-500 Hz) & 0.5-1 Hz per case was 1.7 minutes (IQR: 1.5, 1.8). (**Table 2**)

**Table 2.** Logistic regression models to predict active epileptic spasms.

|  | **AUC** | **Best cut-off value** | **Sensitivity** | **Specificity** | **Accuracy** | **P-values\*** | **Analysis Time (per case)<sup>§</sup>** |
| --- | --- | --- | --- | --- | --- | --- | --- |
| HFOs (80-500 Hz) (/min) - automated + visual verification | 0.98 | 0.51 | 0.91 | 1.00 | 0.93 | < 0.001 | 32.9 min (27.5, 45.2) |
| HFOs (80-500 Hz) (/min) - automated only | 0.96 | 1.13 | 0.83 | 1.00 | 0.87 | < 0.001 | 6.7 min (5.9, 7.1) |
| Ripples (80-250 Hz) (/min) - automated only | 0.95 | 1.09 | 0.83 | 1.00 | 0.87 | < 0.001 | 5.6 min (4.9, 5.9) |
| Fast ripples (250-500 Hz) (/min) - automated only | 0.88 | 0.43 | 0.87 | 0.86 | 0.87 | < 0.001 | 5.2 min (4.9, 5.7) |
| MI HFOs&3-4Hz (automated) | 0.84 | 0.16 | 0.74 | 0.86 | 0.77 | 0.007 | 0.6 min (0.55, 0.65) |
| MI HFOs&0.5-1Hz (automated) | 0.80 | 0.27 | 0.70 | 0.86 | 0.73 | 0.06 | 1.7 min (1.5, 1.8) |
| Hypsarrhythmia (visual analysis) | 0.61 | Presence | 0.22 | 1.00 | 0.40 | 0.09 | 19.9 hrs (18.6, 22.8)<sup>¶</sup> |

\*A Chi-Squared test was used to test the significance of the logistic regression model.
§Values are presented in median (IQR). The computation was performed using a laptop PC with Windows 10 Home (64-bit), Intel(R) Core (TM) i5-7200U (2.50 GHz with 8.00 GB RAM).
¶The presence of hypsarrhythmia was typically determined after the recording was completed. Thus this analysis time indicates the total duration of the recording in each case.

### 3.4. Prediction models based on HFOs rate and MIs of HFOs & SWA

Next, we aimed to predict active epileptic spasms based on the EEG data. Logistic regression models were created using each predictive variable (HFO rates in each frequency band, MI HFOs & 3-4 Hz, MI HFOs & 0.5-1 Hz, and the presence of hypsarrhythmia), from 30 EEGs data from 30 children (if serial EEGs existed, we used the initial studies). (**Table 2**). Based on the small number of the subjects, we did not include other clinical variables such as age, duration of spasms before the EEG, and etiology in the prediction model. In the model incorporating any HFO-related metrics (HFO rate and MIs) performed better than that incorporating the presence of hypsarrhythmia. The prediction model using HFO (80-500 Hz) rate with visual verification exhibited the best area under the receiver operator curve (ROC) value at 0.98, with a sensitivity of 91%, specificity of 100%, and accuracy of 93%.

We then validated the prediction model incorporating the HFO (80-500Hz) rate with visual verification using a different dataset consisted of the second EEG data in 8 children who had serial EEG recordings (two of them had active epileptic spasms). Using the cut-off value of HFO rate ≥ 0.51 (given by the prediction model to give the maximum accuracy), the overall predictive performance was favorable with the sensitivity of 83.3%, specificity of 100%, and accuracy of 87.5%.

### 3.5. Treatment effects on serial EEGs

Two out of eight children who had serial EEG studies responded to medical treatment (adrenocorticotropic hormone, high-dose prednisolone, and/or vigabatrin). The 2 responders had a significant reduction in all three metrics (HFO rate, MI HFOs & 3-4 Hz, and MI HFOs & 0.5-1 Hz) (p < 0.001 with the Wilcoxon signed-rank test, comparing changes in each electrode) (**Figure 2**). However, 6 non-responders showed variable changes (i.e. two had a reduction; two had an increase; two had no statistically significant differences).

### 3.6. Focality of scalp EEG HFOs in relation to neuroimaging findings

There were five patients with focal lesions on neuroimaging studies. We compared the absolute asymmetric indices (abs AI) of HFO (80-500 Hz) rate, MI of HFOs (80-500 Hz) & SWA (3-4 Hz and 0.5-1 Hz) between subjects with a focal lesion on MRI (n = 5) and without (n = 18). Although all the HFO-related values seemed to have higher absolute AI in the group with a focal lesion on MRI compared to that without a focal lesion, none of the group met the statistical significance (**Figure 5**).

**Figure 5:**
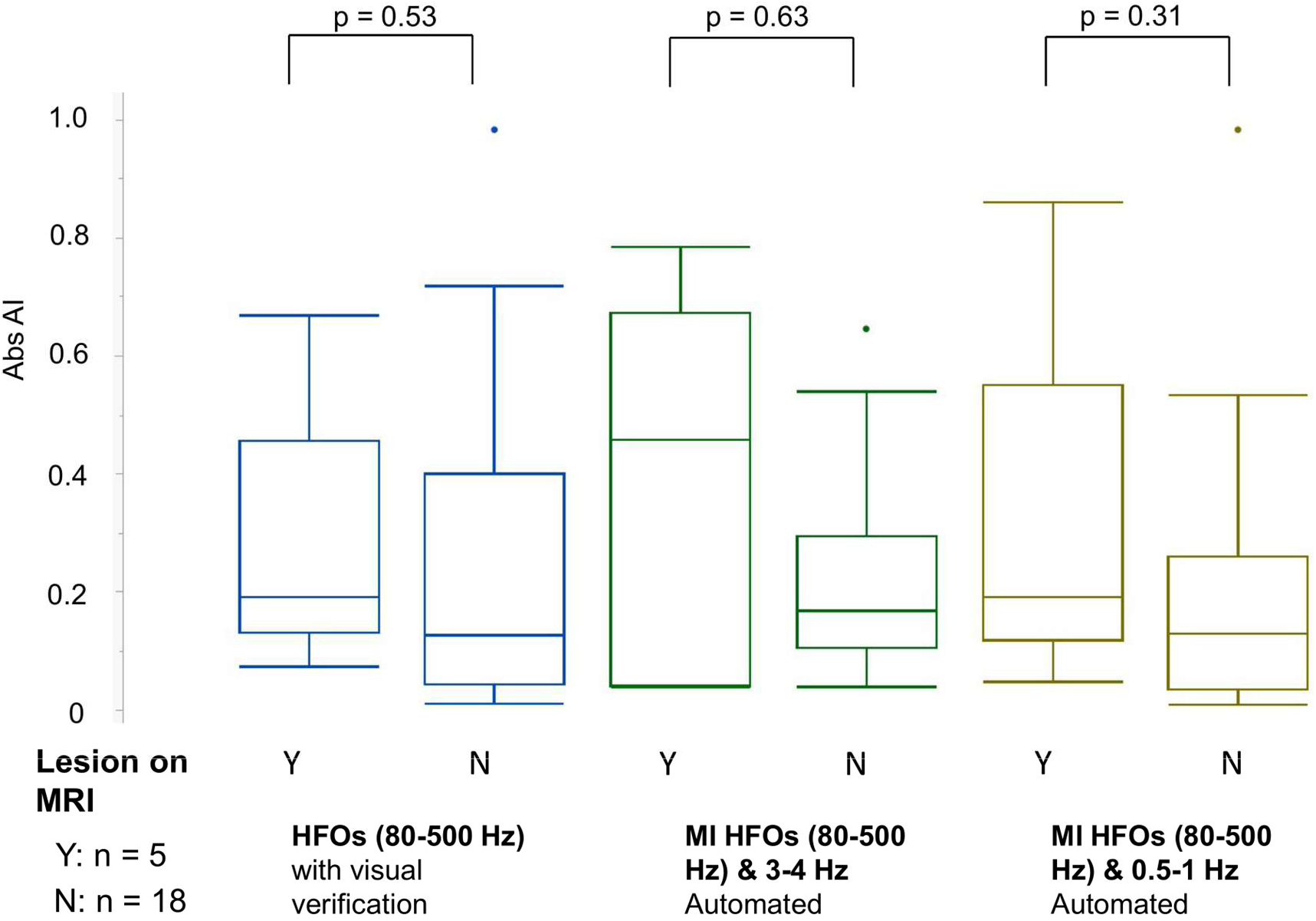
HFOs and neuroimaging correlation. Absolute asymmetric indices (abs AI) of HFO-related values (HFOs with visual verification, MI of HFOs & 3-4 Hz and 0.5-1 Hz) are plotted as a function of the presence (Y) or absence (N) of the MRI lesion. Note all the abs AI looked higher in a group with the presence of the MRI lesion, but none of the group reached statistical significance. AI: asymmetry index. AI = (R-L)/(R+L).

The neuroimaging findings and HFO-related values in each case are summarized in **Table 3**. There are inconsistent findings regarding the side of the lesion and asymmetry of HFO-related values. Especially, patient #29 had right-sided hemimegalencephaly, yet HFO-related values were higher in the left hemisphere, mostly prominently over the left posterior temporal areas (except MIs of HFOs&3-4 Hz SWA being equivocal). The patient underwent right hemispherectomy after the study, and he has been seizure-free over two years.

**Table 3.** HFO-related values and MRI lesion.

| Subject # | MRI Lesion | AI HFOs | AI MI HFOs & 3-4Hz | AI MI HFOs & 0.5-1Hz |
| --- | --- | --- | --- | --- |
| 13 | L hemimegalencephaly s/p hemispherectomy.<br>Persistent epileptic spasms from R hemisphere. | 0.67 | 0.78 | 0.86 |
| 17 | L posterior parietal focal cortical dysplasia | 0.07 | -0.04 | 0.05 |
| 24 | R temporal cortical tuber | 0.24 | 0.46 | 0.24 |
| 25 | R hemispheric focal cortical dysplasia | 0.19 | 0.56 | 0.19 |
| 29 | R hemimegalencephaly | -0.19 | 0.04 | -0.19 |

AI: asymmetry index. $AI = (R-L)/(R+L)$ . Positive numbers indicate higher values on the right side.

## 4. DISCUSSION

We demonstrated that objective analyses of scalp EEG interictal HFO rate and MIs of HFOs and SWA were feasible in children with infantile spasms. High HFO rates and MI values of HFOs & SWA correlated with the presence of active epileptic spasms. After the successful treatments (medications or surgery), the HFO-related values decreased. Such findings were consistent with the prior studies showing active epilepsy correlated with the presence of scalp EEG HFOs in infantile spasms (Kobayashi et al., 2015), benign Rolandic epilepsy (Kobayashi et al., 2011; van Klink et al., 2016), and epilepsy with continuous spikes and waves during slow-wave sleep (CSWS) (Cao et al., 2019; Kobayashi et al., 2010). Although prior studies have reported the presence of scalp EEG gamma band (30-80 Hz) and ripple band (80-250 Hz) activity in infantile spasms (Kobayashi et al., 2004; Kobayashi et al., 2015; Nariai et al., 2017), this is the first study to report that a fast ripple band (250-500 Hz) is seen on scalp EEG in children with infantile spasms. Recently our group reported for the first time that scalp EEG fast ripples could be seen in children with drug-resistant epilepsy with tuberous sclerosis complex (TSC) (Bernardo et al., 2018). Our findings suggest that scalp EEG HFOs including fast ripples (250-500 Hz) can be feasibly analyzed in children with drug-resistant epilepsy.

Importantly, our data indicate HFOs could be a more accurate interictal EEG biomarker of infantile spasms than hypsarrhythmia. The diagnosis of infantile spasms poses a significant challenge to clinicians. The current standard is an EEG test to demonstrate the presence of hypsarrhythmia and epileptic spasms (Lux and Osborne, 2004; Pellock et al., 2010; Wirrell et al., 2015). However, detecting “hypsarrhythmia” in interictal EEG is quite subjective and prone to poor inter-rater reliability (Hussain et al., 2015). In addition, electroencephalographers need an extended EEG study, and typically one recorded during overnight sleep, which may not be feasible in certain geographic regions. More objective and accurate interictal EEG biomarker of infantile spasms, which can be analyzed in a short EEG segment, has been desperately needed. The current study demonstrated that objective analysis of scalp EEG HFOs was quite feasible and could be done in a timely fashion. Although visual verification of the detected HFO events took some time (median 33 minutes per case), the automated detection part was swiftly performed (up to 6.7 minutes per case). Surprisingly, the prediction model incorporating the automated HFO (80-500 Hz) analysis without visual verification performed as well as the model incorporating HFO (80-500 Hz) detection with visual verification (area under the ROC curve: 0.98 vs. 0.96). The false-positive rate of the RMS detector that we used was 13%, which seems tolerable. The substantial advantage of simply using automated detection would be its reproducibility. Visual identification of HFOs requires effort from experienced experts, and inter-rater reliability is suboptimal (Nariai et al., 2018; Spring et al., 2017). Then we would face a similar limitation as in the determination of hypsarrhythmia.

The localization value of scalp EEG HFOs in this population is unclear. In three patients (#13: hemimegalencephaly; #24: TSC; and #25: FCD), HFOs localized to the hemisphere with a presumed epileptogenic MRI lesion. However, one patient (#17: FCD) did not show clear lateralization and another patient (#29: right hemimegalencephaly) had higher HFOs rates contralateral to the MRI lesion (maximal over the left posterior temporal regions). After the surgery, this latter patient became seizure-free over two years. Thus, we believe scalp HFOs, without distinguishing normal and pathological HFOs, could “falsely lateralize,” as is often the case with epileptic spikes. This interpretation is consistent with a recent study demonstrating that scalp HFOs lateralized in the healthier hemisphere in patients with atrophic hemisphere with the epileptogenic lesion, confirmed with the surgical outcome (Kobayashi et al., 2017). A recent study reported that scalp ripples with spike, not ripple without spike localized the epileptogenic zone (Tamilia et al., 2020). Most ripples without spike were observed outside of the epileptogenic zone, thus they speculated that physiological scalp ripples can be detected on scalp EEG. Although our study did not quantify and differentiate HFOs with or without spike, we also observed authentic HFO detection in neurologically normal children, although the detection rate was quite low (0.13/min). Our findings are consistent with their observation, and further study will be needed to quantify and investigate the nature of physiological HFOs detected on scalp EEG.

One of the major limitations of this study was the use of sleep EEG relatively free from artifacts. In order to obtain sleep EEGs, overnight video-EEGs are typically needed. It would be quite challenging to obtain a sleep EEG with a 30-minute routine EEG. As this study’s analysis was solely performed on sleep EEGs, we do not necessarily expect we would get similar results from awake EEGs. To accurately evaluate scalp EEG HFOs on awake EEGs, we would need methods to correct a variety of EEG artifacts, including muscle, movements, and poor impedance (Benar et al., 2010). In this study, we visually excluded “false HFOs” due to such artifacts in the calculation of HFOs rate, but more effort would be needed to analyze awake EEGs. Since we did not analyze the entire EEG for HFOs, we do not know the false-negative rate (how many true HFOs we missed with the detector) and whether selecting different 10 min episodes of SWS would produce the same results. However, with the assumption that there would be abundant HFOs in infantile spasms (Kobayashi et al., 2015), we employed the current approach. The fully-automated calculation of MIs seemed to give us the reasonably accurate estimate of the chance of having active spasms in a timely fashion (median analysis time was 36 seconds in calculating MIs of HFOs & 3-4 Hz SWA). However, we are aware and acknowledge that without verification of the HFOs detection, falsely high MIs from ringing artifact from spikes, electrode or muscle artifact, or spectral leakage of slower oscillations might occur. When fully automated HFOs detection with the capability of event verification is established, we might be able to calculate more accurate MIs between HFOs and SWA. Another limitation is that we included a small number of patients and controls (n=30 in total) in this study. Having a greater number of subjects (n > 100) would have helped create more accurate models, possibly with multivariable regression models to correct variables including age, etiology of epilepsy, and medication effects. In the current study, we validated the model in different EEG data in the same patients. This approach might lack external validity. Ideally, we would have liked more subjects to validate the prediction model in a different set of patients.

Our goal is to establish a robust analytical method to pre-process EEG to remove artifacts so that we can analyze both awake and asleep EEGs. Awake EEGs are much easier to obtain. Having a larger number of subjects and employing completely objective analytic methods on both awake and asleep EEGs will help establish a robust platform to utilize scalp HFOs as a clinically useful EEG biomarker of children with infantile spasms.

## 5. CONCLUSION

Automated analyses of scalp EEG high frequency oscillations (HFOs) and cross-frequency coupling between HFOs and slow-wave activity (SWA) were feasible in children with infantile spasms. Increased rate of HFOs, as well as cross-frequency coupling between HFOs & SWA, can predict active epileptic spasms better than hypsarrhythmia alone.

## 6. ACKNOWLEDGMENTS

The authors have no conflict of interest to disclose. HN is supported by the Susan Spencer Clinical Research Training Fellowship in Epilepsy from the American Academy of Neurology, with funding from the American Epilepsy Society, the American Brain Foundation, and the Epilepsy Foundation. HN also has received research support from the Pediatric Victory Foundation. RS, as well as HN, have received research support from Sudha Neelakantan & Venky Harinarayan Charitable Fund. SAH has received research support from the Epilepsy Therapy Project, the Milken Family Foundation, the Hughes Family Foundation, the Elsie and Isaac Fogelman Endowment, Eisai, Lundbeck, Insys, Zogenix, GW Pharmaceuticals, UCB, and has received honoraria for service on the scientific advisory boards of Questcor, Mallinckrodt, Insys, UCB, and Upsher-Smith, for service as a consultant to Eisai, UCB, GW Pharmaceuticals, Insys, and Mallinckrodt, and for service on the speakers’ bureaus of Mallinckrodt and Greenwich Bioscience. RS serves on scientific advisory boards and speakers bureaus for and has received honoraria and funding for travel from Eisai, Greenwich Biosciences, UCB Pharma, Sunovion, Supernus, Lundbeck Pharma, Liva Nova, and West Therapeutics (advisory only); receives royalties from the publication of Pellock’s Pediatric Neurology (Demos Publishing, 2016) and Epilepsy: Mechanisms, Models, and Translational Perspectives (CRC Press, 2011). RJS is funded by NINDS R01 NS106957, NS33310, NS084017, U54 100064, and Department of Defense EP180003. JYW has received research funding from Novartis, GW Pharmaceutical, NINDS/NIH (R01 NS082649, U01 NS082320, U54 NS092090, U01 NS092595), and Tuberous Sclerosis Alliance, and serves on the scientific advisory boards and speakers’ bureaus for Novartis and GW Pharmaceuticals. The research described was also supported by NIH/National Center for Advancing Translational Science (NCATS) UCLA CTSI Grant Number UL1TR001881. We are indebted to Jason T. Lerner, Joyce H. Matsumoto, Lekha M. Rao, Rajsekar R. Rajaraman, Conrad Szeliga, Maria Garcia Roca, Richard Le, Patrick Wilson, and Kristina Murata for their assistance in the study and sample acquisition.

## Notes

### Competing Interest Statement

The authors have declared no competing interest.

